# Preserved cognitive control with altered ventromedial prefrontal activity in lichen simplex chronicus: an fMRI study during the Multi-Source Interference Task

**DOI:** 10.64898/2026.11.12.766224

**Authors:** L. Valenzuela, J. Valdés, M. Domic-Siede, M. Curi, M. Solis-Avaca, NA. Crossley, F. Zamorano, M. Sandoval, T. Ossandón, V. Medel

**Author notes:** Equally contributing authors (L. Valenzuela, J. Valdés). Shared senior authorship (T. Ossandón, V. Medel).

## Abstract

Lichen simplex chronicus (LSC) is a chronic pruritic skin condition, and one of its central features is the itch-scratch cycle. It has been hypothesized that resisting the urge to scratch, and thereby stopping this cycle, may depend on mechanisms of cognitive control. Here, we used the Multi-Source Interference Task (MSIT) during fMRI to examine whether patients with LSC (n = 9) differ from healthy controls (n = 11) in interference control or in the brain activity that supports it. Our results show that both groups had the expected interference effect, with no statistically significant group differences in behavioral performance or in cingulo-frontal-parietal BOLD activity. Whole-brain analysis revealed a greater interference-related decrease in ventromedial prefrontal cortex activity in patients than in controls. These exploratory findings show distinct patterns of brain activity in LSC while preserving interference control, warranting further investigation in larger samples.

## Introduction

Lichen simplex chronicus (LSC), is a chronic pruritic dermatosis characterized by skin thickening of leathery appearance (lichenified plaques) that result from persistent scratching in a localized area (Ju et al., 2022). The intense itch of LSC disrupts and impairs the quality of life of the patients (Moshkovich et al., 2025). LSC may develop as a primary condition, often associated with psychological or environmental triggers, or arise as secondary to an underlying dermatosis, neuropathy, or systemic disease (Moshkovich et al., 2025). Its pathophysiology is thought to involve neural sensitization and persistent inflammatory signaling (Moshkovich et al., 2025). The itch is thought to be largely non histaminergic mediated by type 2 cytokines that activate C fibers responsible of itch sensing (Ju et al., 2022; Mack & Kim, 2018), and evidence reveals that the lesional skin shows a significant decrease in intraepidermal nerve fibers, consistent with a localized small fiber neuropathy (Sandoval et al., 2022). The core feature of LSC is the itch-scratch cycle, in which scratching disrupts the skin barrier and exacerbates neuroimmune signaling, further intensifying the itch (Moshkovich et al., 2025; Mack & Kim, 2018). Interrupting this cycle is therefore a central therapeutic goal (Ju et al., 2022).

LSC is also highly associated with psychological traits and psychiatric conditions. Patients have nearly twice the odds of a psychiatric diagnosis (Haq et al., 2026), and report elevated anxiety, depression, and perceived stress even in the absence of a psychiatric diagnosis (Altunay et al., 2021). Patients with LSC can also exhibit greater obsessive compulsive traits when compared to healthy controls (Kouri et al., 2016). These psychological features coexist with the behavioral pattern in which scratching can become automatic and reinforced by the immediate itch relief (Tey et al., 2013). Moreover, scratching in the LSC has been reported as particularly pleasurable (Golpanian et al., 2020), potentially contributing to the persistence of the itch-scratch cycle.

These observations raise questions about the cognitive processes involved in regulating scratching in LSC. Although scratching provides immediate relief, repeated scratching conflicts with the longer term goal of preventing further skin damage. Resisting the urge to scratch may therefore draw on mechanisms of cognitive control, which enables goal-directed behavior despite competing response tendencies (Miller & Cohen, 2001; Badre, 2025). Consistent with this possibility, a recent fMRI meta-analysis identified itch-related deactivation in medial frontal and paracentral regions, suggesting that itch processing may interact with neural systems involved in cognitive control (Kim & Day, 2024). However, whether patients with LSC also differ in broader cognitive control processes, such as interference control, remains unknown.

The Multi-Source Interference Task (MSIT) assesses interference control and reliably engages the dorsal anterior cingulate and the cingulate-frontal-parietal network (CFP) (Bush et al., 2003; Deng et al., 2018). Here, we used MSIT-fMRI to examine whether patients with LSC differ from healthy controls in interference control or in the brain activity that supports it, and whether any group differences in brain activity relate to interference effects.

## Materials and methods

### Participants

Thirteen patients with LSC and 11 healthy controls were recruited. The LSC group comprised 8 men and 5 women (M = 46.1, SD = 13.4 years) and the control group 8 men and 3 women (M = 46.0, SD = 13.9 years). Disease duration ranged from 6 to 360 months (M = 84 months, SD = 101 months). Itch intensity was rated from 0 to 10 on the VAS scale (Reich et al., 2012). Median ratings were 8 (range 4-10) for the previous 24 hours, 8 (range 3-10) for the previous week, and 9 (range 3-10) for the previous month of the recruited sample. Quality of life impairment was assessed with DLQI (Finlay & Khan, 1994), with a mean score of 10.7 (SD = 6). Two patients did not undergo MRI scanning, and two other patients were excluded because they did not perform the task as instructed. The final sample comprised 11 controls and 9 patients. The study was approved by the Institutional Ethics Committee of the Pontificia Universidad Católica de Chile (reference number: *160530010*), and all participants provided written informed consent.

### Experimental design

To assess interference control, participants performed the Multi-Source Interference task (Bush et al., 2003) using a blocked design consisting of two conditions, congruent and incongruent. The task consisted of 10 blocks (five per condition) of 24 trials, presented in alternating order, preceded and followed by 30 seconds of rest (fixation on three zeros, ‘000’). On each trial, an array of three digits was presented at the center of a screen, comprising one target digit that differed from two identical distractor digits. Participants were instructed to identify the target digit (1, 2, or 3) while ignoring its spatial position and respond using a response pad with the index (1), middle (2), or ring (3) finger of their dominant hand. In the congruent condition, the target digit appeared in a spatial position matching its numerical identity, and the distractors were zeros (e.g. ‘100’, ‘020’, ‘003’). In the incongruent trials, the spatial position of the target digit was incongruent with the required response, and the distractors were two identical non zero digits (e.g. ‘221’, ‘112’, ‘332’, ‘311’, etc.) (Figure 1). Each stimulus was presented for 750 ms, with 1.82 seconds between stimulus onsets. Participants were instructed to look steadily at the center of the screen presenting the stimuli to avoid artifacts related to eyes and muscle movement.

**Figure 1.**
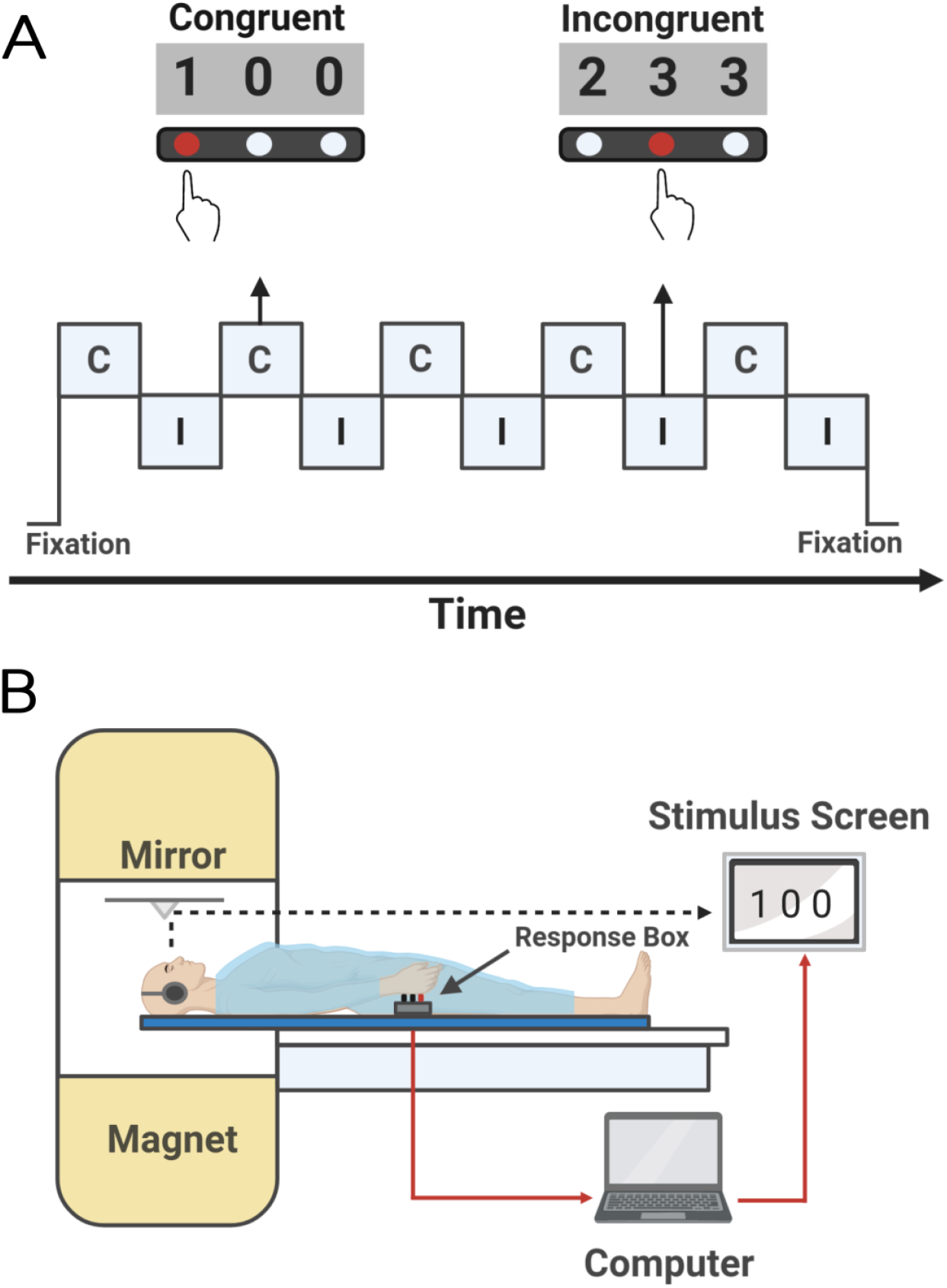
**A)** Block design of the Multi-Source Interference Task (MSIT). **B)** Illustration of the experimental setup during the fMRI acquisition.

### Behavioral and statistical analysis

For each participant and condition, mean reaction time (RT) was computed from correct responses only, and accuracy was calculated as the percentage of correct responses across all trials, with omissions counted as errors. The MSIT interference effect was calculated as the difference between mean incongruent and congruent RTs (incongruent - congruent).

Within-group differences between conditions were tested with Wilcoxon signed-rank tests. Between-group differences in RT and accuracy for each condition, and in the interference effect, were tested with Mann-Whitney U tests. All tests were two tailed, with an alpha significance level of 0.05, and p-values were corrected with Holm’s method across these nine comparisons. Data were analyzed with in-house scripts in Python 3.10 using SciPy (Virtanen et al., 2020) and statsmodels (Seabold & Perktold, 2010).

### Image acquisition and preprocessing

Images were acquired on a Philips Ingenia 3T scanner with a 16-channel head coil (Table 1). Structural images were T1-weighted 3D turbo field echo images, and functional images were T2*-weighted echo-planar images (251 volumes).

**Table 1:** Acquisition parameters of the structural and functional MRI sequences.

| **Acquisition Parameters** | **T1-3D Turbo Field Echo** | **fMRI** |
|---|---|---|
| TR (ms) | 7.8 | 2000 |
| TE (ms) | 3.6 | 35 |
| Matrix (mm) | 240 x 240 | 80 x 80 |
| Field of view (mm) | 240 x 240 x 164 | 220 x 220 x 132 |
| Acquired Resolution (mm<sup>3</sup>) | 1.00 / 1.00 / 1.00 | 2.75 / 2.75 / 3.00 |
| Reconstructed Resolution (mm<sup>3</sup>) | 0.50 / 0.50 / 0.50 | 2.75 / 2.75 / 3.00 |
| Flip angle (º) | 8 | 82 |
| Inversion Time (ms) | 977 | - |
| Bandwidth (Hz) | 191.5 | 35.9 |
| SENSE factor | 2.5 | 1.8 |
| Number of acquisition | 327 | 40 |
| Dynamic scans | - | 251 |
| Acquisition Time | 4 min 8 s | 8 min 27 s |

Functional data were analysed with FEAT (v6.00; FSL version 5.0.8). Preprocessing included motion correction (MCFLIRT), brain extraction (BET), spatial smoothing with a 4 mm FWHM Gaussian kernel, high-pass temporal filtering (cut-off 100 s) and pre-whitening (FILM). First-level statistics were computed in each participant’s native functional space. Functional images were registered to the participant’s T1-weighted image using boundary-based registration (BBR), and the T1-weighted image to the MNI152 2mm template using a 12-degree-of-freedom affine transformation (FLIRT) registration. The two transformations were concatenated, and first-level contrast estimates and their variances were resampled to MNI space for group analysis.

### Image analysis

The model included two explanatory variables (congruent and incongruent blocks), each convolved with a gamma haemodynamic response function (SD = 3 s, mean lag = 6 s). The temporal derivative of each was added as a separate regressor to absorb small timing shifts.

Contrasts weighted only the main regressors (derivatives weighted 0) and estimated congruent and incongruent activity, each relative to the implicit fixation baseline, and their difference. We report the interference effect as incongruent > congruent. Group-level analysis used FLAME 1+2 mixed-effects modelling with automatic outlier de-weighting. The design included one regressor per group (controls, patients) and two variance groups, so that between-subject variance was estimated separately for controls and patients. Contrasts tested each group mean and both group differences. Clusters were defined at Z > 2.3 and corrected at p < 0.05.

To characterize task-related activation in regions typically associated with interference processing during the MSIT, regions of interest (ROIs) were defined using the Schaefer et al. (2018) atlas (400 parcels, 17 networks; MNI152 2 mm). These included the dorsal anterior cingulate/pre-supplementary motor area (bilateral SalVentAttnA medial frontal parcels), and bilateral intraparietal sulcus and dorsolateral prefrontal cortex (ContA parcels), which together form the cingulo-fronto-parietal (CFP) network (Bush & Shin, 2006). Contrast estimates were averaged within each ROI and then across the five ROIs to obtain a single CFP value per participant and condition, expressed as approximate percent signal change. The incongruent > congruent contrast was examined within these regions as a complementary analysis. The within-group differences between conditions were tested with paired t-test and between-group differences with Welch’s t tests (two tailed). Primary statistical inference was performed at the whole-brain level using cluster-based correction. Contrast estimates were also extracted from the cluster identified in the between-group comparison for descriptive analysis. Sensitivity analysis included six motion parameters as confound regressors. Clusters were anatomically labelled with the AAL3 atlas (Rolls et al., 2020) and their network assignment obtained from the Schaefer atlas.

### Brain-behavior relationship

As an exploratory analysis, the incongruent - congruent contrast estimate was averaged across the cluster identified in the whole-brain group analysis and expressed as percentage of signal change. Because the cluster was defined by the group difference, these values were not tested for group differences. RT interference with correct responses was modelled with robust linear regression including the cluster interference BOLD values, group (LSC and HC) and their interaction (HC as reference). Slopes within each group were taken from the same model.

### Meta-analytic decoding

To characterise the functional profile of the group-difference brain cluster, we performed reverse-inference decoding using the Neurosynth database (Yarkoni et al., 2011) with NiMARE (*NeurosynthDecoder*) (Salo et al., 2023). Neurosynth links reported activation coordinates from fMRI literature to the terms used in its text, weighted by term frequency-inverse document frequency (TF-IDF). A study is considered to be associated with a term when its TF-IDF weight exceeds 0.001. To restrict decoding to psychological constructs and avoid anatomical or technical terms, we used 35 cognitive and affective terms derived from the psychological topics of the Neurosynth topic model of de la Vega et al. (2016), plus anxiety and motivation, added due to their relevance as comorbidities for chronic itch and scratching. For each term, we estimated the probability of the term given the activation in the cluster (*P(term|activation)*), assuming a uniform prior of 0.5, so that values above 0.5 indicate a positive association. Associations were tested with chi-square tests and corrected with FDR-BH.

## Results

### Behavior analysis

To test whether a demanding cognitive task affected patients with LSC differently from HC, subjects performed the Multi-Source Interference Task (MSIT). HSC reaction times were slower in incongruent than in congruent trials (Mdn = 803 ms vs 575 ms, W = 0, p = 0.009), with a similar effect observed in LSC (Mdn = 740ms vs 555 ms, W = 0, p = 0.027) (Figure 2A). Consistent with this effect, accuracy was lower in incongruent than in congruent trials in both HC (Mdn = 94.2% vs 100% p = 0.009) and in LSC (Mdn = 93.3% vs 100%, p = 0.047) (Figure 2B). Crucially, there was no statistical difference between the groups in any behavioral measure. Reaction times were similarly modulated by the condition (Congruent U = 67, p = 0.98; Incongruent U = 64, p > 0.99, Figure 2A) as well as accuracy (Congruent U = 60.5, p > 0.99; Incongruent U = 48, p > 0.99, Figure 2B), and the interference effect (Incongruent - Congruent) did not differ between HC and LSC group (HC = 219 ms; LSC = 236 ms; U = 48, p > 0.99, Figure 2C). Thus, subjects showed the expected MSIT behavioral interference effect, with no difference between LSC and HC groups.

**Figure 2.**
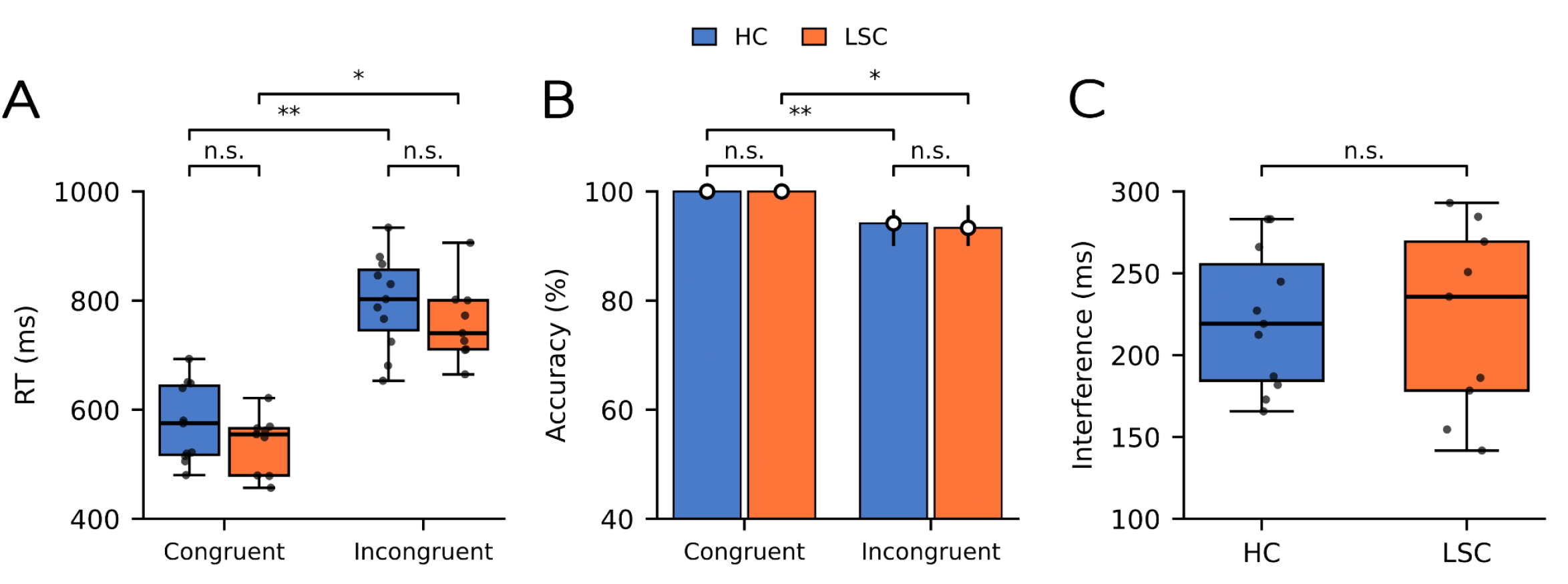
**A)** Reaction time in milliseconds for correct trials only. **B)** Accuracy (median and IQR) in percentage. **C)** Interference effect in milliseconds (Incongruent - Congruent RT). Boxes show median and IQR with dots representing participants (HC n = 11, LSC n = 9). Within-group was done with Wilcoxon tests and between-group comparison was done with Mann-Whitney tests (Holm correction). * p < 0.05, ** p <0.01; n.s., not significant.

### Imaging analysis

Although groups did not differ behaviorally, LSC could differ in the brain activity supporting cognitive interference. We therefore examined the cingulo-fronto-parietal (CFP) network, typically engaged in the MSIT. Both groups robustly engaged the CFP network, with greater BOLD activity in Incongruent than Congruent trials in HC (congruent: Mean = 0.35, SD = 0.25 vs incongruent: Mean = 0.71, SD = 0.22 % signal change, paired t(10) = 7.88, p < 0.001) and in LSC (congruent: Mean = 0.41, SD = 0.22 vs incongruent: Mean = 0.72 SD = 0.22 % signal change, t(8) = 3.60, p = 0.007) (Fig. 3). The groups did not differ in the interference effect (t(12.2) = 0.45, p = 0.66; Fig. 3B) or in either condition (Congruent p > 0.05, Incongruent p > 0.05), and none of the independent ROIs composing the CFP network differed between groups when tested alone (all p > 0.05). Interference is therefore consistently recruiting the CFP network to a similar extent in patients and controls

**Figure 3.**
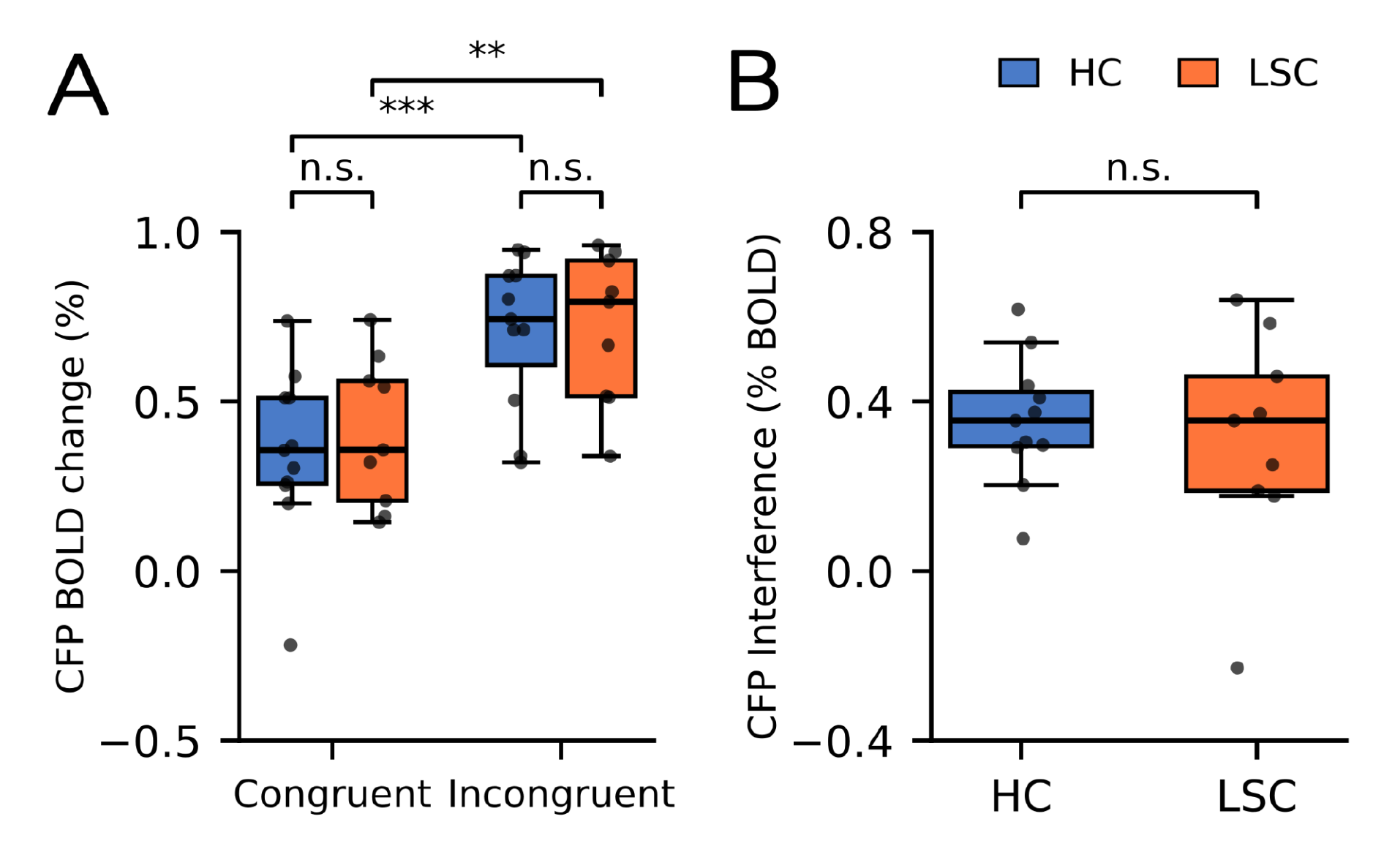
**A)** BOLD signal change by condition averaged across the MSIT ROIs (dACC/pre-SMA, DLPFC, and IPS). **B)** Interference effect (incongruent - congruent) BOLD signal change. HC n = 11, LSC n = 9. Boxes show median and IQR, with dots representing the participants. **p < 0.01, *** p < 0.001, n.s., not significant.

In contrast to specialized interference regions, the whole-brain comparison revealed a significant cluster in the vmPFC, extending into subgenual and pregenual anterior cingulate cortex, where the interference effect (incongruent > congruent) was lower in LSC than in HC (1172 voxels, peak z = 3.80 at MNI 10, 30, −6; cluster-corrected p < 0.001; Table 2; Fig. 4A). Its cortical portion lay within the Default A network of the Schaefer atlas (400 parcel, 17 network). The effect remained significant when six motion parameters were included as regressors (732 voxels, peak z = 3.65, MNI 6, 34, 8; cluster-corrected p < 0.001; Supplementary Fig. S2), with its cortical portion also within the Default A network, and mean framewise displacement did not differ between groups (p > 0.05). Two smaller clusters, in the right supramarginal/inferior parietal and right posterior temporal cortex, showed larger interference related increases in HC than in LSC but did not survive the inclusion of motion regressors. To illustrate the direction of this difference, we extracted values from the vmPFC cluster, and found that in LSC, activity decreased from congruent to incongruent trials (interference effect Mean = −0.31 % signal change, SD = 0.20), whereas no change was detected in HC (Mean = 0.00 % signal change, SD = 0.16; Fig. 4B-C). Consistently, within-group maps showed a significant decrease in these regions in patients only, whereas controls showed decreases elsewhere in the Default A network (Supplementary Fig. S1). Therefore, the only robust group difference is distributed not in regions involved in cognitive interference, but in a ventromedial default-mode region that deactivates with interference only in LSC patients.

**Figure 4.**
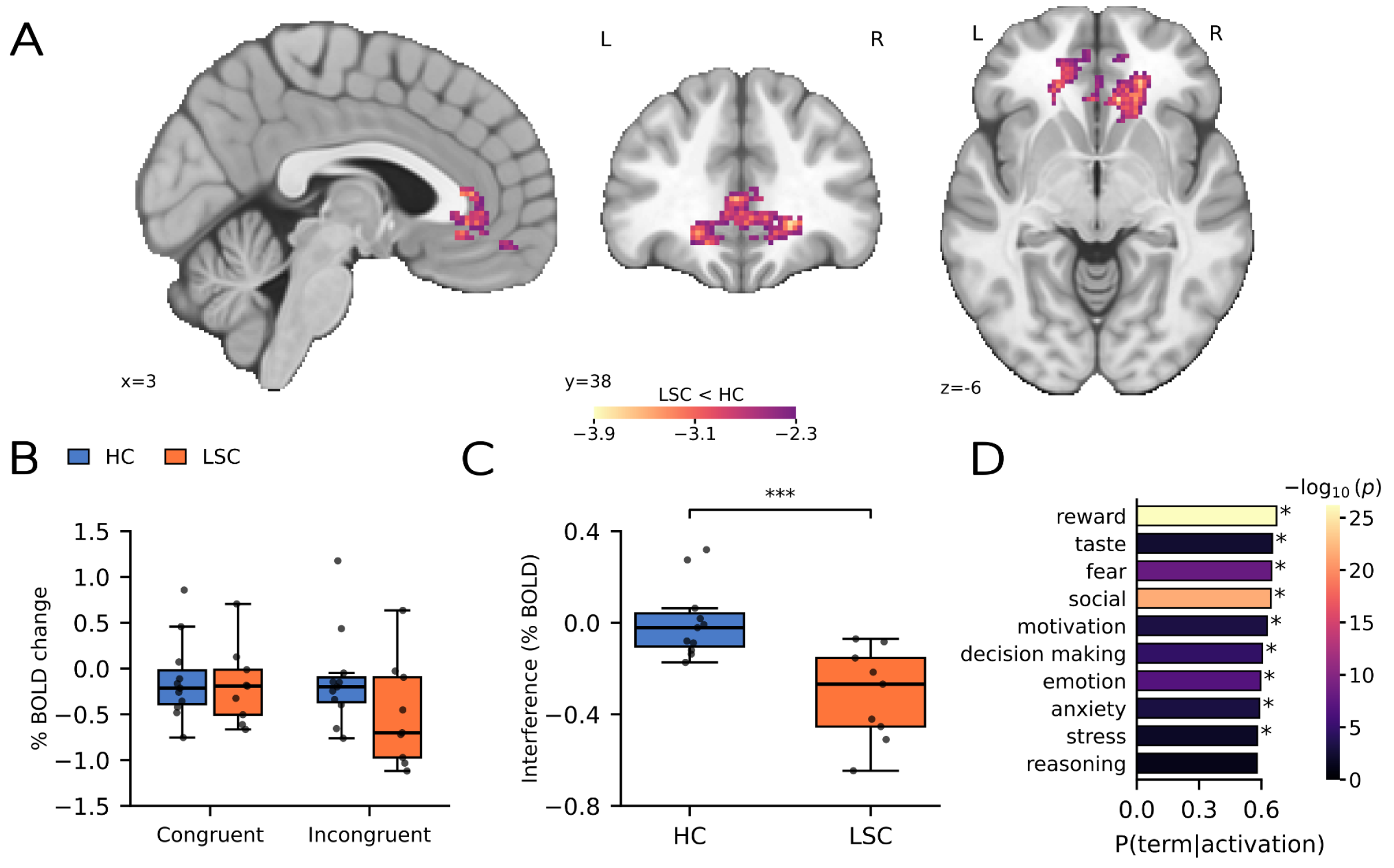
**A)** Whole-brain cluster in which the interference effect (incongruent > congruent) was lower in LSC than in HC (FLAME 1+2, cluster-forming Z > 2.3, cluster-corrected p < 0.05, peak at MNI 10, 30, −6) **B, C)** Values extracted from the cluster shown for descriptive purposes. **B)** % BOLD change for each condition relative to the resting fixation baseline and, **C)** interference effect (incongruent - congruent); asterisks indicate the whole-brain cluster level effect *** p < 0.001, **D)** Neurosynth decoding of the significant cluster showing the 10 terms with the highest P(term|activation), colour indicates −log10(p), * p < 0.05 corrected with FDR. Boxes show median and IQR.

**Table 2:**
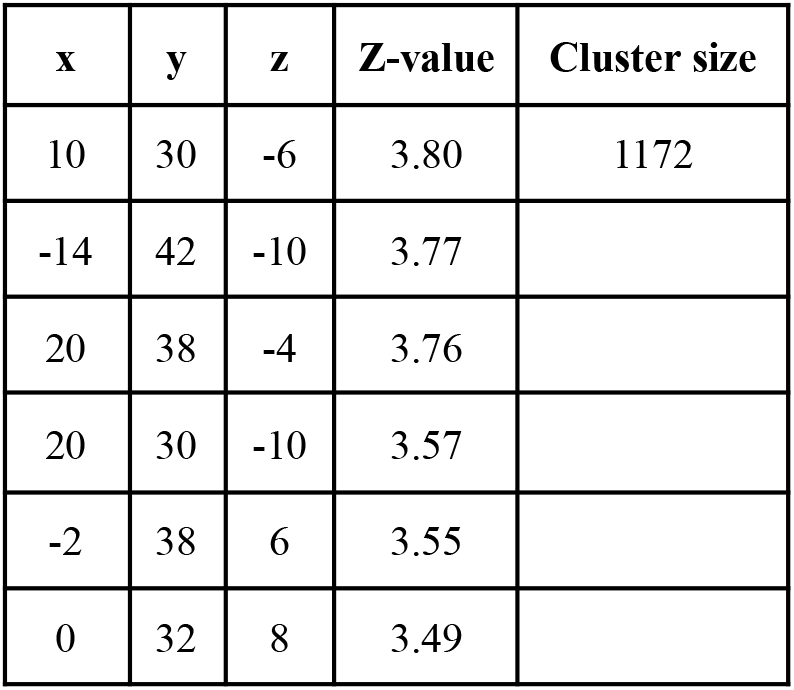
Local maxima of the vmPFC cluster showing a group difference in the interference effect (HC > LSC for incongruent > congruent; FLAME 1+2, cluster forming Z > 2.3, cluster-corrected p < 0.001). Coordinates are in MNI152 space. Of the cluster’s voxels, 28% fell in AAL3 Frontal_Med_Orb (vmPFC), 15% in ACC_pre, 11% in ACC_sub, and 36% in white matter. Within the Schaefer atlas, 99% of its cortical voxels lay within the Default A network.

| **x** | **y** | **z** | **Z-value** | **Cluster size** |
|---|---|---|---|---|
| 10 | 30 | -6 | 3.80 | 1172 |
| -14 | 42 | -10 | 3.77 |  |
| 20 | 38 | -4 | 3.76 |  |
| 20 | 30 | -10 | 3.57 |  |
| -2 | 38 | 6 | 3.55 |  |
| 0 | 32 | 8 | 3.49 |  |

### Brain and behavior relationship

To further explore whether the vmPFC cluster was related to behavior, we explored the relation between RT interference and cluster interference and did not found statistical relation neither in HC (b = 130.9 ms per % signal change, SE = 104.4 p = 0.21) nor LSC (b = −8.6, SE = 92.5, p = 0.93). No statistically significant difference was found when comparing the brain-behavior slopes between groups (robust regression, group x cluster interaction b = −139.5 ms per % signal change, p = 0.32), indicating that the vmPFC cluster difference was not statistically accompanied by a difference in how this activity relates to behavioral interference.

### Meta-analytic decoding

Finally, to characterize the functions associated with the vmPFC cluster, we performed a meta-analytic decoding. Of the studies that appeared in the Neurosynth database, 1733 reported activation within the significant cluster. The cluster was most strongly associated with reward (P(term|activation) = 0.67), taste (0.65), fear (0.65), and social (0.65), followed by motivation, decision making, emotion, anxiety, stress and pain (0.57 to 0.63 all p < 0.05 FDR corrected, Fig. 4D), where values above 0.5 (the prior) indicate a positive association. Importantly, cognitive control terms were not associated and were at or below chance level (conflict 0.46; inhibition 0.48). This meta-analytic result suggests that the BOLD group differences arose in a region associated with key cognitive terms involved in valuation, reward and affective systems rather than cognitive control or interference.

## Discussion

In this exploratory study we asked whether lichen simplex chronicus patients showed an impairment in the behavioral and neural responses associated with interference control in comparison with healthy controls. Our results indicate that both groups exhibited the expected interference effect elicited by the MSIT (Fig. 2), with no statistically significant difference between groups in behavioral performance (all p > 0.05). In line with cognitive interference, both groups showed greater BOLD activity during incongruent than congruent trials in the dACC/pre-SMA, DLPFC, and intraparietal sulcus (Fig. 3), regions typically engaged by the MSIT (Bush & Shin, 2006; Deng et al., 2018). These regions, analyzed together here as the cingulo-frontal-parietal (CFP) network (Bush & Shin, 2008), also showed no significant group differences in their responses to interference either when tested together or individually (Fig. 3B). These findings do not support impaired interference control in LSC under the conditions tested.

Skin and mind modulate each other in many chronic dermatoses (Cameron et al., 2024). This is consistent with LSC, where patients show a distinctive psychological profile marked by negative affect, worry and harm avoidance, obsessive-compulsive traits, and scratching that becomes automatic activating a pleasured experience. Previous fMRI studies have linked these components to the brain. Indeed, stress-induced itch relates to hippocampal and subcortical activation in patients with LSC or prurigo nodularis (Kim et al., 2016), Furthermore, scratching an itch deactivates the vmPFC and ACC in close relation to its pleasurability in healthy controls (Papoiu et al., 2013), and reward regions became overactive during the itch-scratch cycle in chronic itch patients (Mochizuki et al., 2015). Interestingly, control systems may limit the itch, as suggested by the association between greater frontoparietal connectivity and smaller increases in evoked itch in atopic dermatitis, suggesting possible top-down regulation (Desbordes et al., 2015). Moreover, a recent meta-analysis also proposed a possible role for cognitive control in regulating itch-related urges, while emphasizing the need for direct experimental investigation (Kim & Day, 2024). Our study complements this literature by assessing interference resolution in LSC, while leaving open whether difficulties emerge specifically when resisting the urge to scratch, or its relation with itch intensity symptoms.

The only significant group difference appeared in the whole-brain analysis which identified a greater interference-related decrease in vmPFC BOLD activity in LSC than in controls (Fig. 4B). Within the patients, activity was significantly lower during incongruent than congruent trials, whereas controls showed no significant decrease in this region, although they exhibited decreases elsewhere in the default mode network (Supplementary Fig. S1, Table S1). Decreased activity in default mode nodes are commonly reported during externally focused tasks (Menon, 2023), and conflict-related decreases in these regions have been reported during the MSIT (Wojciechowski et al., 2024), a suppression proposed to depend on salience network engagement and its interplay with control-related nodes (Menon, 2011; Menon, 2023). The meta-analytic decoding associated the vmPFC cluster map with affective and valuation related terms, consistent with accounts of this region involvement in self-related processes and affective meaning (Roy et al., 2012; Hiser & Koenigs, 2018). These associations provide context but do not identify the psychological processes underlying the group difference. One speculative possibility is that itch-related affective or self-focused processing competes with task demands in LSC, and that the greater interference-related vmPFC decrease could reflect attenuation of this processing. Whether this modulation helps sustain task performance remains unknown. Future studies assessing itch-related symptoms across task conditions, including itch intensity and the urge to scratch, and examining their associations with vmPFC responses and performance could evaluate this account. Examining whether these relationships change following symptom relief could further clarify its relevance to the itch-scratch cycle.

Our study has several limitations to be noted. First, the sample size limits statistical power and the precision of the effect estimates, as such, small or moderate differences cannot be ruled out. Second, the exploratory vmPFC finding requires replication in larger samples, and although psychological and clinical measures were collected, their associations with behavioral performance and brain activity were not examined. Thus, whether the observed difference is related to psychological symptoms or other clinical characteristics of LSC, such as itch-intensity, remains unclear.

In conclusion, patients with LSC showed no detectable group differences in behavioral performance or CFP responses to interference, but exhibited a greater interference-related decrease in vmPFC BOLD activity. This exploratory finding requires replication in larger samples and further investigation of its relationship with itch, scratching and psychological symptoms that clarify its potential relevance to the pathophysiological mechanisms of the lichen simplex chronicus.

## Acknowledgments

We thank the participants and their families for their invaluable contributions. We also thank Martin Irani, Nicolás Gravel and Roberto García for their valuable methodological comments. We acknowledge the open-source tools and data that we used for this research. V.M. is supported by ANID/FONDECYT de Iniciación 11251578, ANID/FONDECYT Exploración 13240170, and AARG-25-1490306 from the Alzheimer’s Association. T.O. is supported by ANID/FONDECYT Regular 1230383. NC, TO and MD-S are supported by Centro de Interés Nacional IINARA, CIN250068. JV is supported by Chilean National Center for Artificial Intelligence CENIA FB210017, Basal ANID.The content is solely the authors’ responsibility and does not represent the official views of these institutions.

## Supplementary material

**Figure S1.**
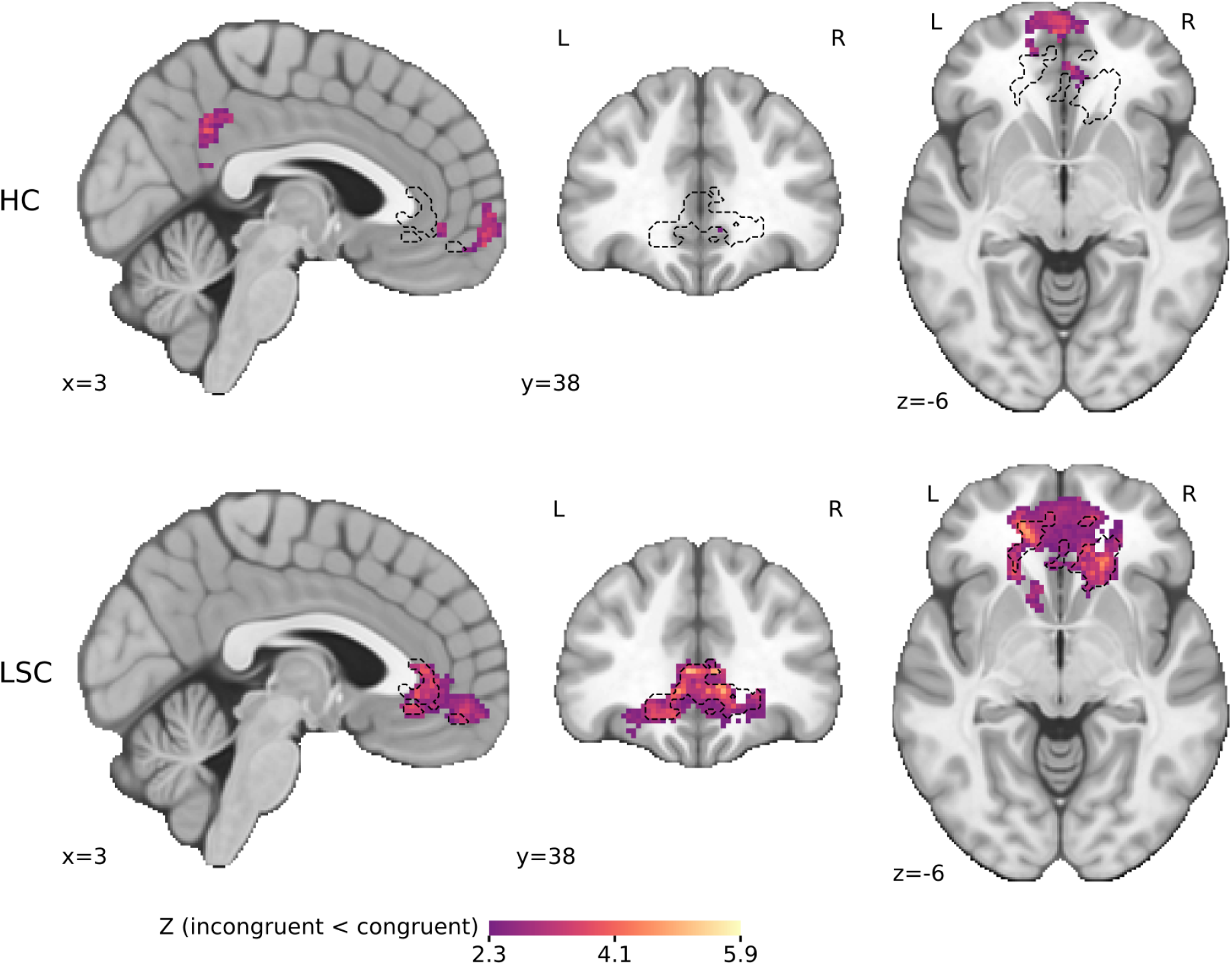
Interference related decreases (incongruent < congruent) within HC (top) and LSC (bottom) (FLAME 1+2, cluster-corrected p < 0.05), predominantly within the Default A network in both groups. Dashed lines show the vmPFC significant between-group difference.

**Figure S2.**
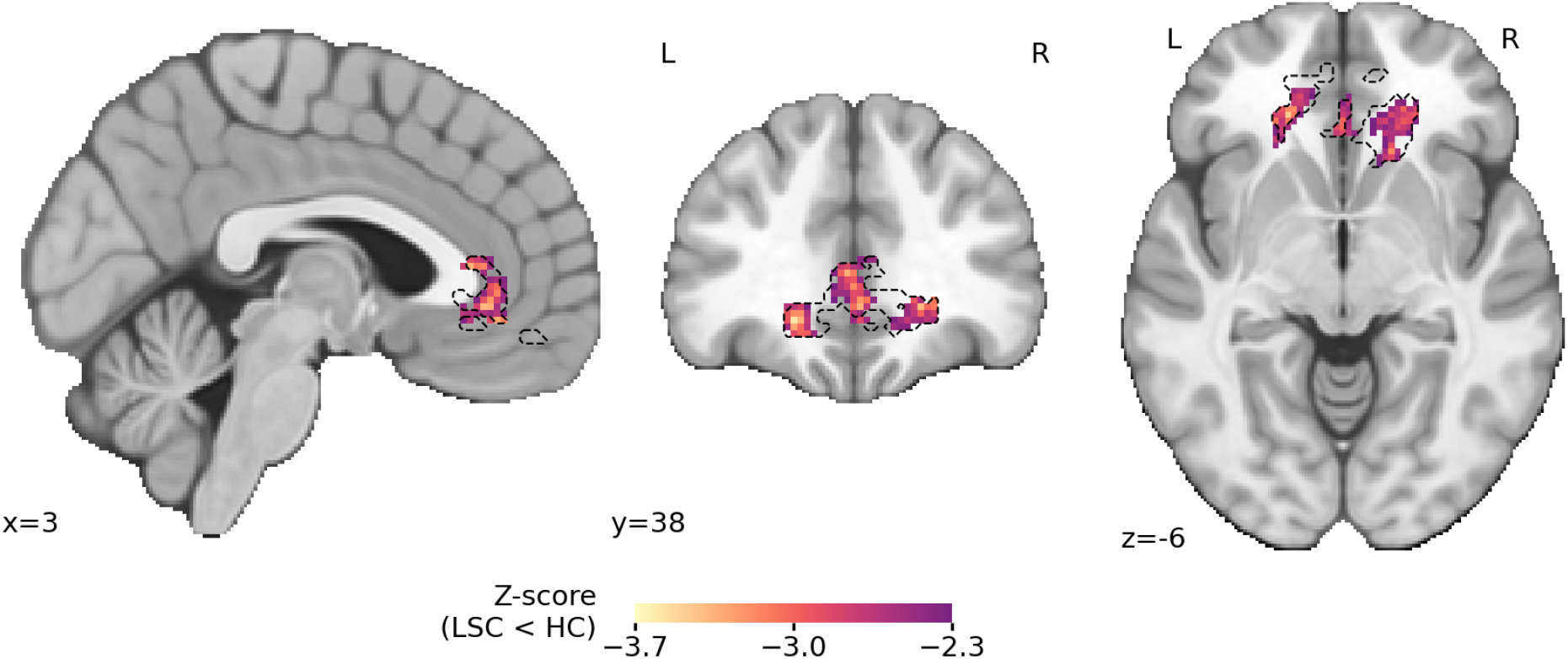
Sensitivity analysis showing the group difference in the interference effect (incongruent - congruent) with LSC < HC, after including the six motion parameters as regressors estimated with MCFLIRT. Dashed lines show the cluster described in the main model.

**Table S1.**
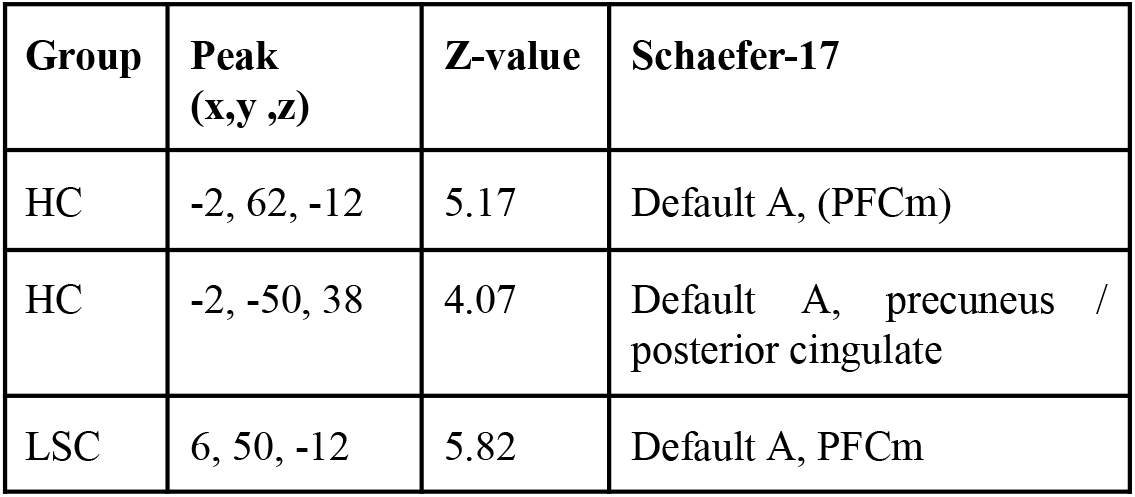
Peak of the clusters showing interference related decreases (incongruent < congruent) within each group (cluster forming Z > 2.3, cluster-corrected p < 0.05). Coordinates in MNI152, regions from the Schaefer 400-parcel, 17-network atlas.

