## Supplementary Materials for "Preserved cognitive control with altered ventromedial prefrontal activity in lichen simplex chronicus: an fMRI study during the Multi-Source Interference Task"

### Supplementary material

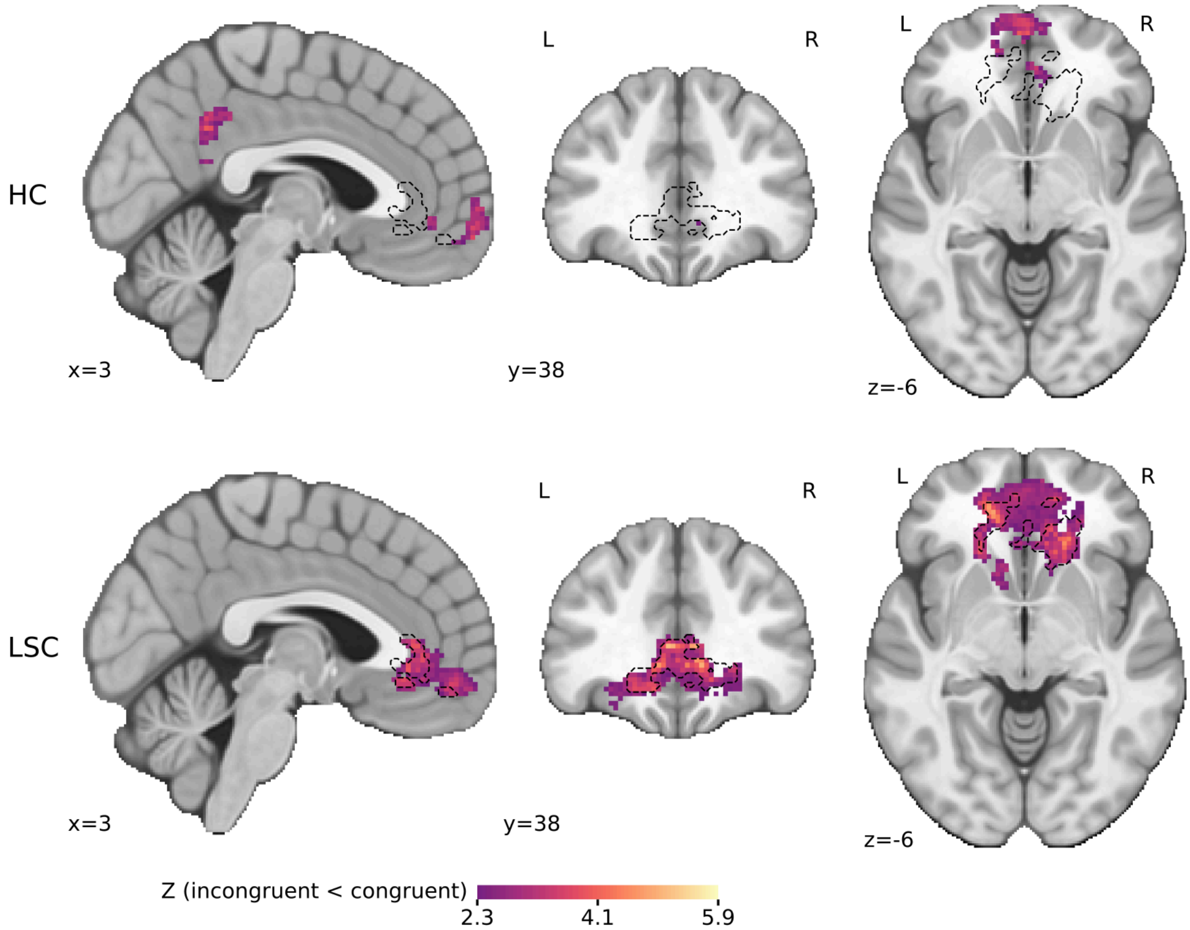


**Figure S1.** Interference related decreases (incongruent < congruent) within HC (top) and LSC (bottom) (FLAME 1+2, cluster-corrected $p < 0.05$), predominantly within the Default A network in both groups. Dashed lines show the vmPFC significant between-group difference.

| Group | Peak (x,y ,z) | Z-value | Schaefer-17 |
|---|---|---|---|
| HC | -2, 62, -12 | 5.17 | Default A, (PFCm) |
| HC | -2, -50, 38 | 4.07 | Default A, precuneus / posterior cingulate |
| LSC | 6, 50, -12 | 5.82 | Default A, PFCm |

**Table S1.** Peak of the clusters showing interference related decreases (incongruent < congruent) within each group (cluster forming $Z > 2.3$, cluster-corrected $p < 0.05$). Coordinates in MNI152, regions from the Schaefer 400-parcel, 17-network atlas.

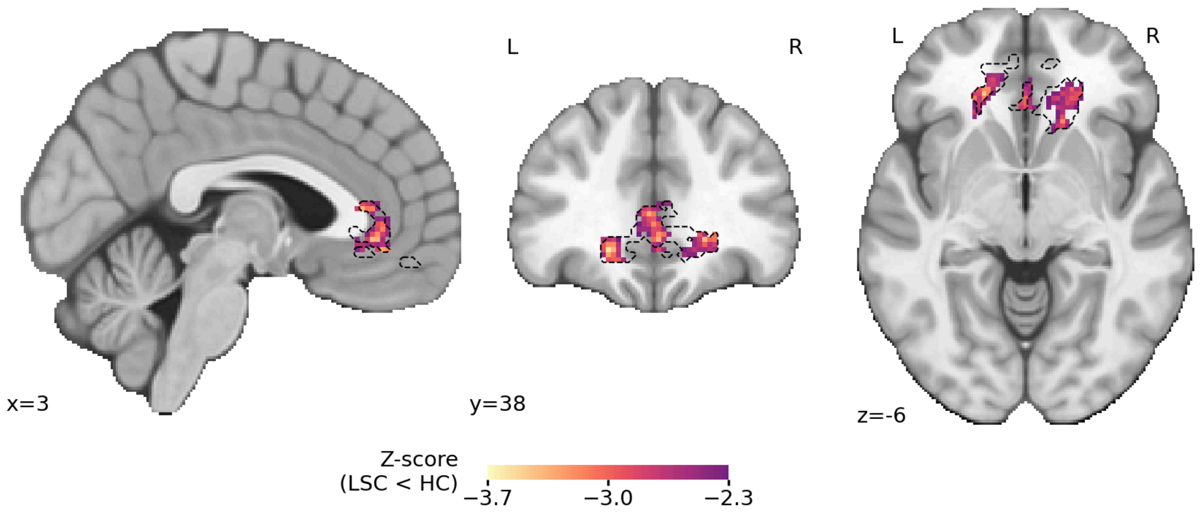


**Figure S2.** Sensitivity analysis showing the group difference in the interference effect (incongruent - congruent) with LSC < HC, after including the six motion parameters as regressors estimated with MCFLIRT. Dashed lines show the cluster described in the main model.
